# Leveraging agent-based models and deep reinforcement learning to predict taxis in cell migration: Insights from barotaxis

**DOI:** 10.1101/2025.04.11.648312

**Authors:** Daniel Camacho-Gomez, Raffaele Sentiero, Maurizio Ventre, Jose Manuel Garcia-Aznar

## Abstract

We present a novel computational framework that combines Agent-Based Modeling (ABM) with Reinforcement Learning (RL) using the Double Deep Q-Network (DDQN) algorithm to determine cellular behavior in response to environmental signals. We showcase its potential by modeling how pressure gradients direct cell migration in confined environments, a phenomenon known as barotaxis. By integrating RL, the model allows cells to learn and adapt their migration behavior based on sensed pressure gradients, capturing the dynamic, environment-dependent nature of cell behavior. We validate the framework using real microfluidic devices and experimental data, demonstrating the model’s ability to predict barotactic migration. Thus, this approach introduces a novel direction for modeling how cells sense and transduce environmental cues into biological behaviors.

## 1. Introduction

Cell migration is a fundamental biological process involved in development, immune responses, tissue repair, and disease progression [1]. Among the factors that influence this complex behavior, external cues play a crucial role in guiding cells as they navigate their environment. Depending on the nature of these signals, cells exhibit directional migration in response to chemical gradients (chemotaxis), surface-bound cues (haptotaxis), stiffness gradients (durotaxis), and geometrical features (topotaxis) [2]. In addition, pressure gradients represent another type of external signal that can guide cell migration through a process known as barotaxis, which has gained increasing attention in recent years [3]. Barotaxis was first proved in *in vitro* experiments using microfluidic devices, which showed that cells migrating in confined asymmetric hydraulic environments tend to move toward the path with the least hydraulic resistance [4]. As cells migrate, they displace fluid, generating pressure gradients based on the hydraulic resistance of their surroundings [5]. These pressure gradients, influenced by the asymmetry of the environment, affect decision-making during migration and guide cell movement [6, 7].

The influence of pressure gradients on cell migration becomes particularly significant in the context of cancer metastasis, a hallmark of cancer [8] and a leading cause of cancer-related mortality [9]. During metastasis, cells must escape the primary tumor and navigate through the tumor microenvironment (TME) [10], a complex and dynamic space characterized by profound alterations in biochemical composition, cellular interactions, metabolic state, and mechanical properties [11]. These mechanical alterations include increased matrix stiffness [12, 13], compression [14], solid stress [15], and elevated interstitial fluid pressure [16]. As a result, the TME becomes highly confined and dynamic, influencing the presence of pressure gradients that can affect cell migration as they move during metastasis. In this regard, [17] studied the migration of cancer cells in asymmetric hydraulic environments, highlighting the preferential migration of cancer cells toward paths with lower hydraulic resistance. Therefore, gaining insight into the mechanisms that drive barotaxis could be crucial for predicting cancer cell metastasis and developing new strategies to disrupt this invasive process.

Computational approaches have become essential in modern biological research due to their ability to model complex biological processes and predict outcomes. Among the various types of models in biology, Agent-Based Models (ABMs) are wide-adopted computational tools that have been widely employed to study cell migration [18–22]. Thus, ABMs allow for the simulation of individual cells and, consequently, investigate the impact of microenvironmental cues on cell behavior [23–25]. Nonetheless, while ABMs provide valuable insights, they need refined calibration techniques to accurately replicate experimental observations.

Machine learning (ML) models are emerging as powerful tools to adapt computational models to data variability, enhancing physics-based models’ ability to capture intricate patterns. When combined with ABMs, ML techniques have the potential to further advance the modeling of cellular behaviors in complex biological systems [26]. Specifically, reinforcement learning (RL) offers powerful frameworks to model complex decision-making processes in dynamic environments [27]. RL enables agents to learn optimal strategies by interacting with the environment, making it highly effective for real-time decision-making tasks. Despite the synergy between ABMs and RL, which has proven to be promising in other fields [28–30] since its introduction by [31], it has yet to be fully applied in biology.

In this work, we aim to predict barotactic cell migration in confined microenvironments through computational modeling. In particular, we present an ABM that integrates an ML model based on RL to predict the influence of pressure gradients on cell migration direction in confined microchannels. To achieve this, we first compute the pressure gradients within the microchannels using Computational Fluid Dynamics (CFD). Then, we simulate cell migration with the ABM, considering the temporal and dynamic variation in the migration direction influenced by the pressure gradients. In this regard, we reproduce the mechanosensing process of cells by adding observation points on the cell membrane where it senses fluid pressure. This sensed pressure is then passed into a Neural Network (NN) that determines the direction of migration. The neural network subsequently adjusts the migration direction based on the pressure gradients sensed by the cell throughout the migration process. To make the cell learn barotactic cell migration decisions, we apply the RL approach, specifically the Double Deep Q-Network (DDQN) algorithm [32]. Finally, we train the model using test geometries and validate it with a real microfluidic geometry designed to study barotaxis from [17], showing agreement between experimental observations and *in silico* predictions. Thus, this novel approach represents a first step toward building an intelligent *in silico* cell that reproduces how cells transduce external cues from the environment into migration behaviors.

## 2. Results

### 2.1 Computational framework for barotactic cell migration

The proposed computational model has three main parts: the environment, the ABM, and the RL algorithm. The environment’s geometry corresponds to the microfluidic device’s migration chamber, where the *in vitro* experiments are conducted [17]. Within this geometry, we perform a CFD simulation to obtain the pressure field *P* (***x***), which serves as the environmental cue guiding cell migration (Fig. 1**A**). Then, we simulate cell migration within this environment using an ABM (Fig. 1**B**). Here, we assume that the temporal variation in the migration direction ***e***(*t*) is influenced by pressure gradients to replicate barotactic migration. To determine how cell migration is affected by these pressure gradients, we employ a NN trained using an RL approach based on DDQN (Fig. 1**C**). In our model, the agent observes the fluid pressure at equidistant points around the cell surface, replicating the cell’s mechanosensing. This sensed pressure is then passed into the NN, which outputs the probabilities of migration towards discrete action points evenly distributed around the surface. This information is used to update the migration direction at each time step (Δ*t*) throughout the ABM simulation.

**Fig. 1.**
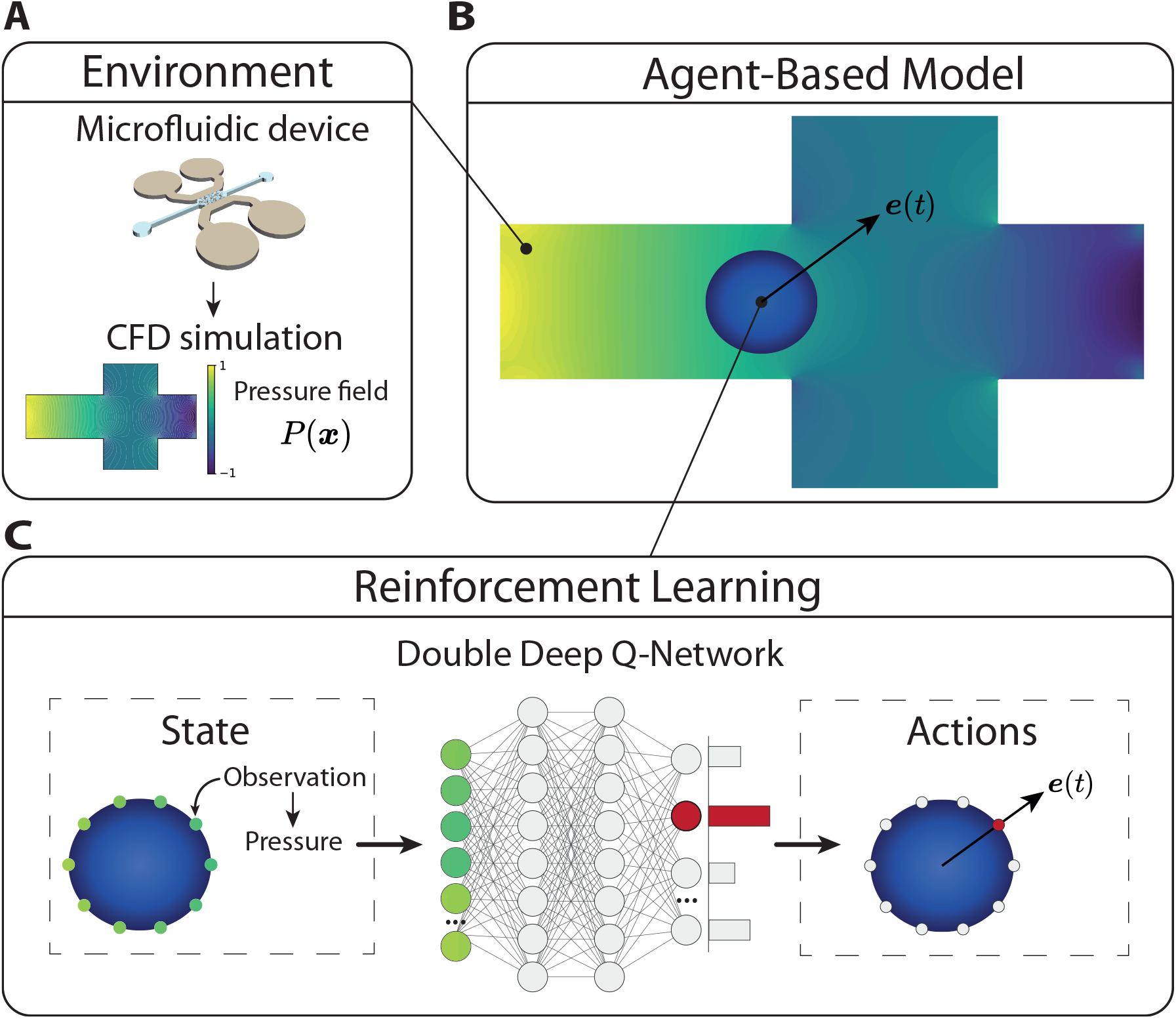
Overview of the computational model. **A** The environment consists of the pressure gradients within the microfluidic device, where *in vitro* experiments are conducted. **B** The ABM simulates the migrating cell, considering the influence of pressure gradients on the migration direction ***e***(*t*). **C** The migration direction ***e***(*t*) is determined through an NN trained using an RL approach based on DDQN. To do that, the NN receives the sensed pressure from the cell and outputs the probabilities of migration toward different directions. Finally, this new migration direction is passed to the ABM, with the process repeated at each time step (Δ*t*) throughout the simulation.

### 2.2 Learning barotactic cell migration behavior

First, we train our model to enable the cell to sense pressure stimuli and respond accordingly by regulating its migration direction, thereby learning barotactic cell migration behavior. To achieve this, the cell (agent) is trained in three different geometries, each featuring a bifurcation point with a three-channel intersection. In each geometry, the outlet boundary condition is placed at different locations after the bifurcation point: the straight channel, the top channel, and the bottom channel. These boundary condition locations are designed to create varying pressure gradients, helping the agent learn how to sense and respond to these gradients by migrating in different directions. In each case, the goal position, ***x***_*g*_, which influences the reward function, is located at the outlet boundary. We simulate the migrating cell in each of the three geometries and minimize the resulting loss using the DDQN method. Consequently, the reward across all training geometries is represented in each episode, yielding a mean reward of 0.9999 after 7,543 episodes (Fig. 2**A**). As a result, the cell successfully learns to migrate in response to pressure gradients in these three training geometries, moving toward the higher pressure gradient (Fig. 2**B**). Consequently, as the cell migrates, we observe a decrease in the mean pressure surrounding it (Fig. 2**C**). The parameters of the DDQN model, along with the parameters of the ABM model, can be found in Table 1.

**Table 1.** Parameters of the computational model.

| Hyperparameter | Value/Description |
| --- | --- |
| Batch size | 512 (number of transitions sampled from the replay buffer) |
| Discount factor ($\gamma$) | 0.1 (importance of future rewards) |
| Epsilon ($\epsilon$) | 0.9, decays to 0.0001 over 1000 episodes |
| Learning rate | $1 \times 10^{-4}$ (for the AdamW optimizer) |
| Soft update rate ($\tau$) | 0.005 (rate at which the target network is updated) |
| Number of observations ($N_o$) | 72 (NN input) |
| Number of actions ($N_a$) | 16 (NN output) |
| Time step ($\Delta t$) | 10 min |
| Cell velocity ($v_c$) | 0.2 $\mu\text{m}/\text{min}$ (mean velocity of the cell from [17]) |
| Radius of the cell ($R_c$) | 10 $\mu\text{m}$ |
| Radius of the cell nucleus ($R_N$) | 4 $\mu\text{m}$ |

**Fig. 2.**
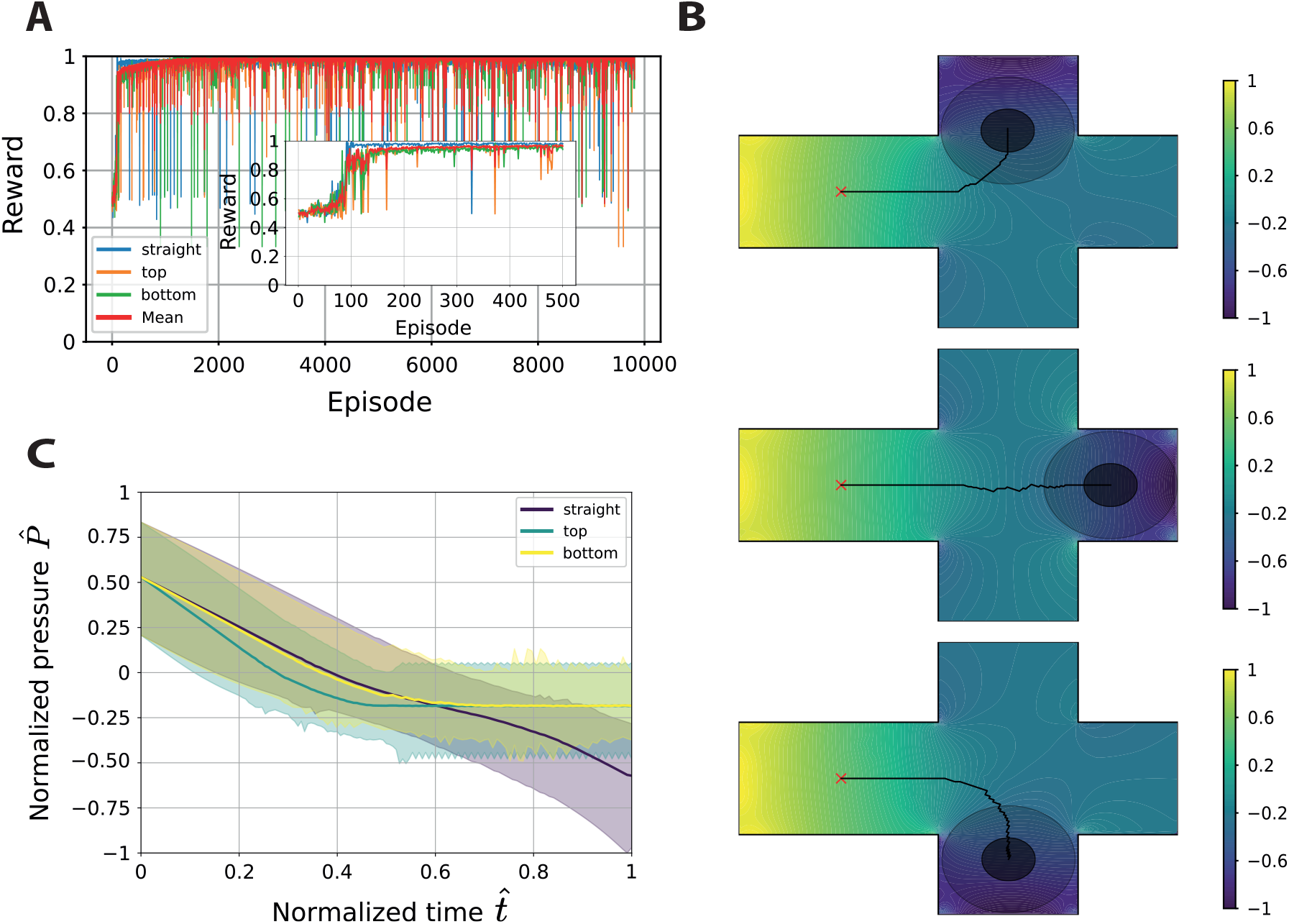
Training results for barotactic migration. **A** Reward across episodes in the straight, top, and bottom geometries, with a zoomed-in detail of the first 500 episodes, reaching a mean value of 0.9999 after 7,543 episodes. **B** Trajectories of the cell across the top, straight, and bottom geometries after the training process. **C** Mean normalized pressure around the cell in each geometry, with shaded regions indicating the minimum and maximum values.

### 2.3 Predicting in vitro barotactic cell migration

To validate our model, we test its capacity to predict barotactic cell migration in three different real microfluidic devices. In particular, we employ the dead-end, twisted, and tortuous microdevices from previous work [17] (Fig. 3). The dead-end microdevice serves as a control device for barotaxis since it maximizes the hydraulic resistance difference between both paths, maximizing the pressure gradient through the top channel. Consequently, 77% of the cells in the *in vitro* experiments migrate through top open-path channel, showing a directed migration bias (Fig. 3**A**, top). Our computational model successfully reproduces barotactic migration, with the cell following the higher pressure gradients to move through the top channel (Fig. 3**A**, bottom left, and Supplementary Video 1). Thus, we can analyze the pressure sensed by the cell throughout its migration (Fig. 3**A**, bottom middle). This chart reveals the asymmetrical pressure distribution around the cell and how it progressively decreases over time as the cell moves in the direction of the pressure gradient. In fact, we also compute the mean pressure around the cell throughout its migration, highlighting the cell’s tendency to move from regions of higher pressure to lower pressure (Fig. 3**A**, bottom right).

**Fig. 3.**
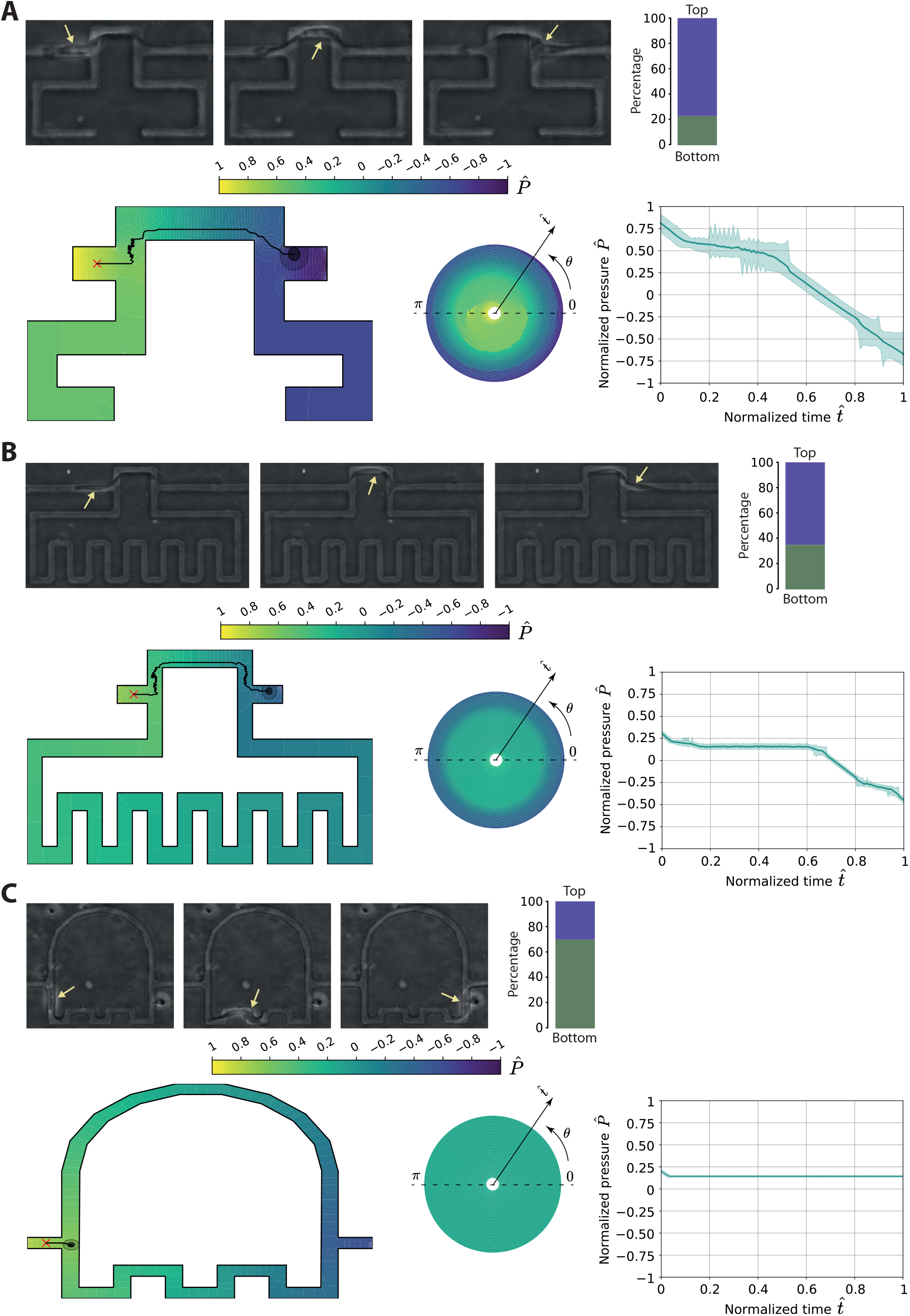
Reproducing *in vitro* barotactic migration. **A** Dead-end microdevice. **B** Twisted microdevice. **C** Tortuous microdevice. For each microdevice, the top panel shows snapshots of the cell migration experiments and the percentage of cells migrating in each channel from [17]. The bottom left panel presents the predicted trajectories. The bottom middle panel displays a donut chart representing the pressure around the cell, where each concentric ring corresponds to the pressure at a given time. The chart includes 50 rings, with the smallest ring representing the initial condition and the largest ring representing the final time. The bottom right panel illustrates the mean normalized pressure around the cell, with shaded regions indicating the minimum and maximum values.

In the case of the twisted microdevice, *in vitro* data show that around 65% of cells migrate towards the top path (Fig. 3**B**, top). Here, the computational model also replicates this cell migration behavior, despite a slight difference in pressure gradients between the bifurcating channels (Fig. 3**B**, bottom left, and Supplementary Video 2). As a result, the pressure distribution around the cell over time is smoother compared to the dead-end microdevice (Fig. 3**B**, bottom middle), and the mean pressure profile during migration remains more consistent (Fig. 3**B**, bottom right). Regarding the tortuous microdevice, approximately 70% of cells are inclined to migrate toward the bottom tortuous path (Fig. 3**C**, top). However, there is almost no pressure difference between the bifurcating paths, so our computational model does not predict any barotactic cell decision at the bifurcation point (Fig. 3**C**, bottom left). Indeed, the pressure distribution around the cell at the bifurcation point is constant, remaining unchanged as the cell does not move from this point (Fig. 3**C**, bottom middle). Finally, the mean pressure sensed by the cell decreases initially as the cell moves from the starting point to the bifurcation point, and then remains constant, with no differences between mean, maximum, and minimum pressure values, indicating a uniform distribution around the cell. Therefore, these results suggest that barotaxis does not explain the preferential cell migration toward the bottom tortuous path, and other mechanisms might be contributing to this migratory behavior.

## 3. Discussion

We presented a novel computational framework to reproduce how cells sense and transduce environmental signals into biological behaviors. To this end, we developed a model that couples ABM with RL using the DDQN algorithm. This integration represents a conceptual advancement in modeling cellular behavior. Rather than relying on predefined parameters, static rules, or heuristics, the RL component enables the cell to learn from its microenvironment and adapt its behavior based on sensed signals. To demonstrate its potential, we simulated how pressure gradients direct cell migration in confined environments, a phenomenon known as barotaxis [4]. To this end, we obtained the pressure field in the confined environment using CFD, which provides cues that the cell transduces into migration behavior. Then, we simulated cell migration with a lattice-free ABM, in which the direction of migration is regulated by the environmental pressure gradients. To simulate how cells sense and respond to these varying pressure gradients, we incorporated an ML model. This model consists of an NN trained through DDQN that receives the pressure distribution around the cell and determines the migration direction. This combination provides a dynamic and adaptable approach to simulating complex biological processes, such as how cells transduce environmental cues into cell behavior. Additionally, it enables the cell to make decisions based on real-time feedback from environmental cues, such as pressure gradients, in contrast to more traditional approaches where behavior might be pre-programmed or governed by simplistic rules. Therefore, it offers a promising framework for better understanding and predicting complex cell behavior in dynamic environments.

To validate our model, we applied our framework to real microfluidic devices and tested its performance against experimental *in vitro* data from the study of [17]. This facilitated the assessment of the model’s ability to accurately predict cell migration behavior in experiments, demonstrating its ability to generalize across different geometries and configurations, while reinforcing the robustness and applicability of the approach. Thus, we reproduced barotactic cell migration in the dead-end microdevice, identifying the environmental cues that the cell transduced into biological migration behavior. Interestingly, we observed that the cells oscillate at the bifurcation points, with direction changes that enable them to correctly sense the environmental cues and mechanotransduce this information into migration decisions, as seen in the oscillations of the pressure in Fig. 3**A**, Supplementary Videos 1 and 2 and in experimental observations [17]. Furthermore, we also predicted barotactic cell migration in the twisted microdevice, demonstrating the model’s sensitivity in reproducing barotactic migration despite, slight differences in pressure gradients between bifurcating channels.

However, we did not predict the migration trajectories in the tortuous microdevice. In this microdevice, there are no differences in pressure between the channels at the bifurcation point. Nevertheless, experimental results show a preferential migration toward the bottom tortuous path. Thus, our model indicates that barotaxis is not the driving mechanism guiding cell migration toward this bottom path, suggesting that other factors may be influencing this behavior, as hypothesized in [17]. Possibly, other migration mechanisms may be involved in this directed migration [33], such as the geometric characteristics of the bottom channel, which might provide mechanical support that allows cells to attach to it rather than the curved channel [2]. Therefore, our model serves to predict and identify whether barotaxis is the underlying mechanism driving cell migration observed in experimental data. The potential of this approach is not limited to barotactic cell migration. This computational framework can be applied to model directed cell migration in response to other factors, such as chemical gradients (chemotaxis), surface-bound cues (haptotaxis), stiffness gradients (durotaxis), and geometrical features (topotaxis) [2]. More importantly, in real biological systems, different environmental stimuli often coexist, competing to guide cell migration and potentially outcompeting each other [34].

Therefore, our approach has the potential to contribute to a more comprehensive understanding of the directional decision-making processes in cell migration.

Nonetheless, it is important to acknowledge that our approach relies on certain simplifications, which must be thoroughly justified. The most significant of these relates to the assumption of a time-independent pressure field within the microdevices. In this work, we simulated the fluid flow within the geometry to obtain a static pressure field. However, the actual mechanism involves the cell migrating through the confined channel, pushing the fluid and generating a dynamic pressure gradient as it moves. This effect could be captured by simulating the pressure field resulting from a moving object representing the cell in the CFD simulation. In this case, at each time step, the CFD simulation would be performed with a moving object representing the cell’s movement in the ABM. This would enable a fully coupled, real-time simulation between the CFD and the ABM. However, the computational cost of real-time CFD simulations during the training process was prohibitively high. Therefore, we opted to simulate the fluid flow once to characterize the microfluidic device and obtain a representative pressure field.v

Another simplification is that we do not explicitly represent cell deformation. We assumed the cell to be non-deformable since our focus was on reproducing barotactic cell behavior rather than modeling the exact cell shape. To account for cell-wall interactions, we considered the interactions between the wall and the cell nucleus. In this way, we simplified the cell deformation calculation by assuming that the cytoplasm of the cell is deformable and would not oppose resistance, while the nucleus of the cell is non-deformable and resists deformation when it touches the wall. This is consistent with experimental measurements showing that the cell nucleus is stiffer than the cytoplasm [35–37]. Through this approximation, we were able to simulate cell trajectories with a deformable cell, allowing us to focus on the actual trajectories and cell behavior. In this regard, our simulations show that the cell deforms and migrates close to the walls to minimize the traveled distance. This can be observed in Fig. 3**A** and Fig. **B** (bottom left), where the cell deforms to migrate along the lower wall in the middle section and then moves to the top wall at the outlet. This migration pattern aligns with the *in vitro* experiments, where cells migrate along the predicted trajectories, attaching to the corresponding walls. Through this study, we demonstrate that RL can be a powerful tool for reproducing cellular behaviors in ABMs. By leveraging RL, cells can dynamically determine their behavior by transducing external cues in complex environments, reducing reliance on heuristic or rule-based models. Therefore, this approach has the potential to enhance our understanding of how environmental signals influence biological processes, contributing to more accurate models of cell behavior.

## 4. Methods

### 4.1 Environment

The cell’s environment is defined by the pressure field within the microfluidic device. To obtain this pressure field in the microchannels, we perform a Computational Fluid Dynamics (CFD) simulation by means of the Finite Volume Method (FVM) in Ansys Fluent 2023R2. To solve the fluid flow in the microdevice, we use the material properties of liquid water and set an inlet boundary condition with a uniform velocity profile of 0.2 μm*/*min, which corresponds to the velocity at which the cell pushes the fluid in experiments [17]. The outlet boundary condition was specified as an atmospheric pressure outlet. We assume laminar flow with a low Reynolds number due to the low velocity magnitude and small characteristic lengths of the microdevices. The solvers employed include a pressure-velocity coupled scheme with a third-order MUSCL scheme for momentum and a second-order scheme for pressure.

### 4.2 Agent-based model

The lattice-free center-based ABM simulates the migrating cell within the microdevice. In this model, we calculate the position of the cell ***x***_*c*_(*t*) as:

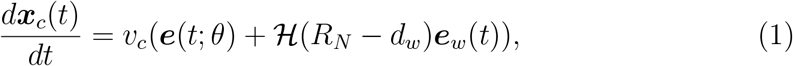

with *v*_*c*_ the magnitude of the velocity, ***e***(*t*; *θ*) the unit direction vector of migration, is the ℋ Heaviside step function, *R*_*N*_ the radius of the cell nucleus, *d*_*w*_ the shortest distance from the cell’s position to the wall, and ***e***_*w*_(*t*) the unit direction vector resulting from the interaction of the cell with the walls. Here, we focus on reproducing pressure-driven cell migration decisions, therefore, we consider a constant velocity magnitude and a variable cell migration direction. The magnitude of the velocity is obtained from the mean velocity of (*v*_*c*_ = 0.2 μm*/*min) measured in the confined experiments in [17]. The direction of migration ***e***(*t*; *θ*) is approximated by a NN with weights *θ* that takes the local pressure values that the cell senses at each position. Thus, the description of the temporal evolution of ***e***(*t*; *θ*), based on the pressure gradients, replicates barotactic cell migration. The cell-wall interaction is triggered when the cell’s nucleus contacts the wall. This interaction is modeled using the Heaviside step function ℋ (*R*_*N*_ −*d*_*w*_), which becomes active when the distance *d*_*w*_ becomes less than or equal to *R*_*N*_. In this case, the unit direction vector ***e***_*w*_(*t*)) opposes the cell’s motion, pointing opposite to the vector from the cell center to the collision point, thereby preventing the cell from moving through the wall.

### 4.3 Double Deep Q-Network

In this study, we employ a Q-learning approach to determine the cell’s direction of migration (action *a*) based on the pressure values sensed by the cell (observation *ô*_*t*_ at a particular time *t*) from the pressure field (state *s*) using OpenAI’s Gymnasium library [38]. Q-learning is a reinforcement learning algorithm that aims to estimate the optimal action-value function, *Q*(*ô, a*), which represents the expected cumulative reward of taking an action *a* based on the observations *ô* from the state *s*, and following an optimal policy thereafter [27]. Specifically, we use Double Deep Q-Network (DDQN) algorithm [32], where a NN approximates the Q-function, *Q*(*ô, a*; *θ*), with *θ* representing the NN’s weights. This model mitigates the overestimation bias in action-value predictions from the standard Deep Q-learning by separating the action selection and evaluation processes. DDQN uses two distinct networks: the policy network, which selects actions, and the target network, which evaluates the selected actions, thereby improving stability and learning performance.

The Q-value for a given observation-action pair (*ô*_*t*_, *a*) is approximated as:

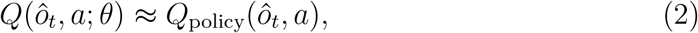

where *ô*_*t*_ is the observation from the state *s* made at time *t*. In our model, the agent observes the fluid pressure values at *N*_*o*_ equidistant points around the surface, replicating the cell’s mechanosensing. To ensure consistency, these observed pressure values are normalized to the range [− 1, 1] using min-max scaling, where the minimum observed pressure maps to − 1 and the maximum to 1. Similarly, the agent’s actions correspond to *N*_*a*_ equidistant discrete points around the surface, so the migration direction ***e***(*t*; *θ*) derives from the unit direction vector pointing from the agent’s center to the activated action. Thus, *N*_*o*_ and *N*_*a*_ control the spatial resolution of environmental sensing and action selection, respectively, influencing how finely the agent perceives pressure cues and determines its migration direction.

The target Q-value is computed as follows, using the target network and the Double Q-Learning update.

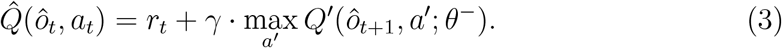

Here, *r*_*t*_ is the reward received after taking action *a*_*t*_ in observation *ô*_*t*_, *γ* is the discount factor that weights future rewards, and *Q*^′^(*ô*_*t*+1_, *a*^′^; *θ*^−^) is the *Q*-value output of the target network for the next observation and a given action *a*^′^ that maximizes the Q-value. The reward *r*_*t*_ is calculated using the following function:

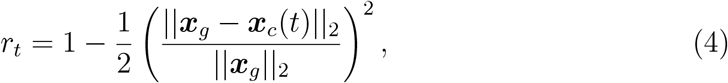

where ***x***_*g*_ represents the goal position and ***x***_*c*_(*t*) is the cell’s position. This reward function encourages the cell to move closer to the goal by increasing the reward as the distance decreases, with larger distances penalized more heavily to promote efficient movement.

To balance exploration and exploitation during training, the agent follows an *ϵ*-greedy policy for action selection. At each time step, it chooses either an action *a*_*t*_ based on the policy network and a random action:

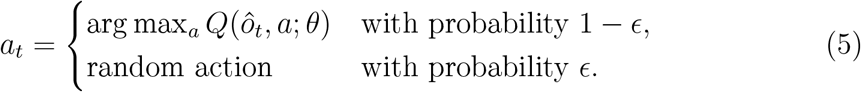

As training progresses, *ϵ* decays to encourage greater exploitation of the learned policy.

We use experience replay to break the correlation between consecutive experiences by storing transitions (*ô*_*t*_, *a*_*t*_, *r*_*t*_, *ô*_*t*+1_) in a replay buffer. During training, random mini-batches of transitions are sampled from the buffer to update the policy network. This improves learning stability by reducing the variance of updates.

The optimization of the policy network is performed by minimizing the Huber loss, a robust loss function for regression tasks, between the Q-values predicted by the policy network and the target Q-values computed using the target network:

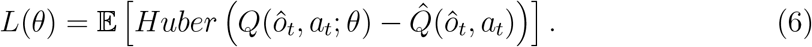

Finally, the target network’s weights are periodically updated based on the policy network’s weights using the soft update rule:

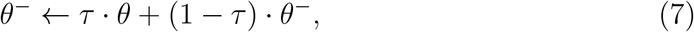

with *θ* the weights of the policy network, *θ*^−^ the weights of the target network, *τ* the soft update rate.

## 5. Acknowledgements

This research is part of a project that has received funding from the European Research Council (ERC) under the European Union’s Horizon 2020 research and innovation programme (ICoMICS grant agreement No 101018587).

